# Effects of radiofrequency electromagnetic radiation (RF-EMF) on honey bee queen development and mating success

**DOI:** 10.1101/434142

**Authors:** Richard Odemer, Franziska Odemer

## Abstract

Mobile phones can be found almost everywhere across the globe, upholding a direct point-to-point connection between the device and the broadcast tower. The emission of radiofrequency electromagnetic radiation (RF-EMF) puts the surrounding environment inevitably into contact with this pollutant. We have therefore exposed honey bee queen larvae to the radiation of a common mobile phone device (GSM) during all stages of their pre-adult development including pupation. After 14 days of exposure, hatching of adult queens was assessed and mating success after further 11 days, respectively. Moreover, full colonies were established of five of the untreated and four of the treated queens to contrast population dynamics. We found that mobile phone radiation had significantly reduced the hatching ratio but not the mating success. If treated queens were successfully mated, colony development was not adversely affected. We provide evidence that RF-EMF only acts detrimental within the sensitivity of pupal development, once succeeded this point, no further impairment has manifested in adulthood. Our results are discussed against the background of long-lasting consequences for colony performance and the possible implication on periodic colony losses.

**HIGHLIGHTS:**

- Chronic RF-EMF exposure significantly reduced hatching of honey bee queens
- Mortalities occurred during pupation, not at the larval stages
- Mating success was not adversely affected by the irradiation
- After the exposure, surviving queens were able to establish intact colonies

**GRAPHICAL ABSTRACT:** 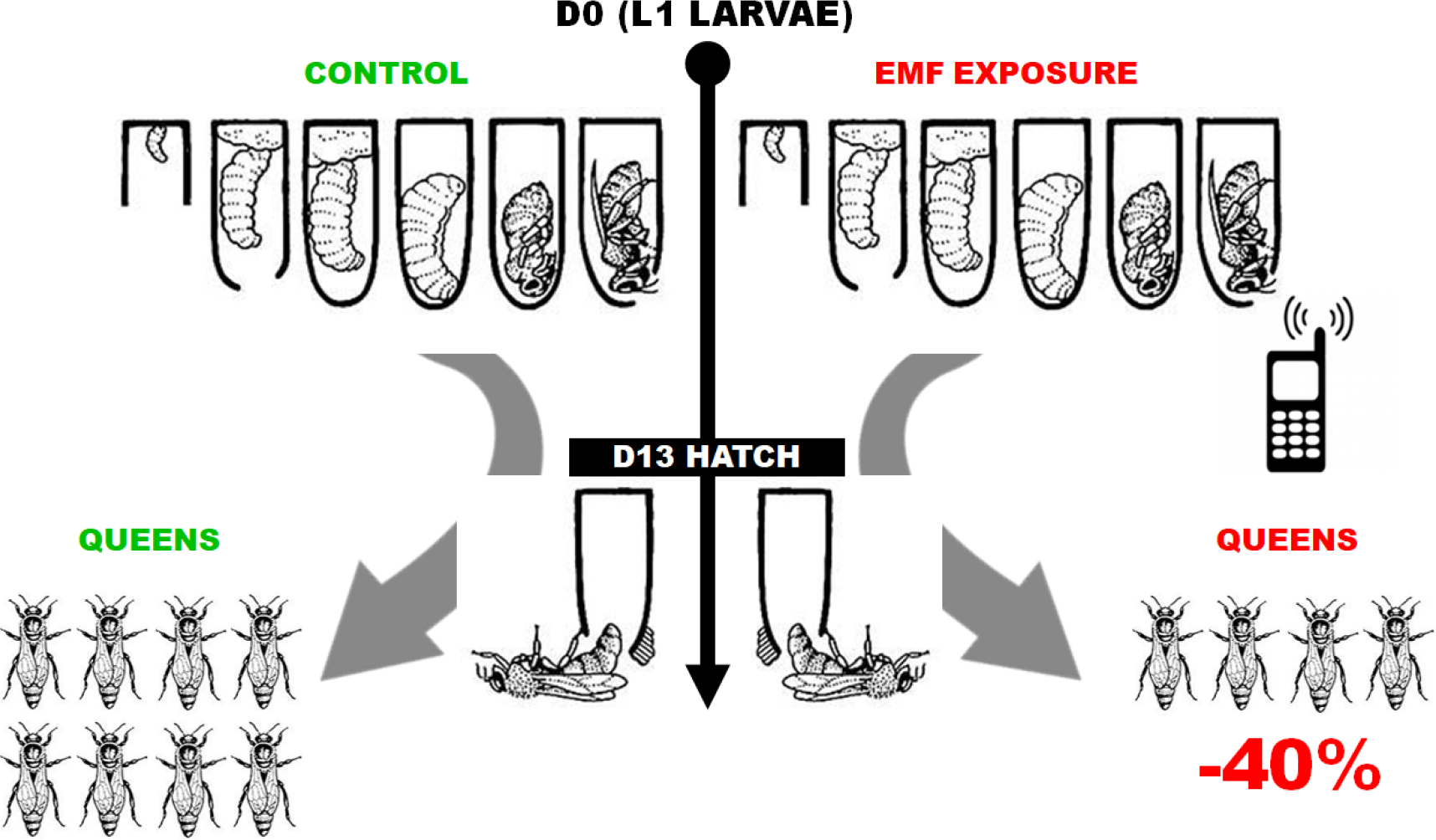

## 1 INTRODUCTION

The modern world turns around technological achievements and it is simply not possible to imagine our everyday life without them. With an estimated 6.9 billion subscriptions globally, mobile phone devices such as smart phones have established their position in our society (WHO, 2014). In many countries, cell phones are important tools not only for communication but also for bank transfers, newscast, social media and numerous other conveniences with an increasing tendency. Provided that this market will be further growing in the future, concerns are rising about the emission of radiofrequency electromagnetic fields (RF-EMF) from these devices and their broadcasting network, i.e. antennas and base stations, perceived as environmental pollution (Balmori 2015).

Radiofrequency waves are electromagnetic fields, and unlike ionizing radiation such as X-rays or gamma rays, they can neither break chemical bonds nor cause ionization in the living tissue (Genuis & Lipp 2012). They are usually ranging from 30 kHz-300 GHz with cell phones operating mainly between 800 MHz and 3 GHz, pulsed at low frequencies (Hardell 2017). As a consequence, they are often strictly forbidden in medical facilities and on airplanes, as the radiofrequency signals may interfere with certain electro-medical devices and navigation systems.

In the last decade field and laboratory studies have furthermore demonstrated that RF-EMF exposure is of ecological relevance. The radiation may have an impact on surrounding flora as well as vertebrate and invertebrate organisms (Cucurachi et al. 2013). Effects have manifested in different ways and some of them are a cause of concern. A large scale monitoring study (> 10 years) revealed that in trees, a closer range to phone masts resulted in significant damages in the side facing the mast in contrast to the opposite side (Waldmann-Selsam et al. 2016) whereas Roux et al. (2006, 2008) found exposed tomato plants to show similar consequences when wounded, trimmed or burnt. In chicken eggs, Batellier et al. 2008 found an increased mortality when exposed to cell phone radiation over the entire incubation period. Very similar to previous study results from Bastide et al. (2001) and Grigoryev (2003), this developmental stage seems to be particularly vulnerable for non-thermal radiation. A proportional relationship between the intensity of the electromagnetic field and the negative effects, however, could not be established (Batellier et al. 2008).

In fruit flies, reproduction and reproductive organs were also significantly affected by mobile phone radiation (Panagopoulos et al. 2004, Panagopoulos 2012) unlike to the findings of Weisbrot et al. (2003) where a beneficial effect on the reproductive success was reported. In their study, the number of offspring increased by up to 50 % compared to control, demonstrating controversial outcomes. Studies in insects have shown that reproduction cycles and change of generations are quick, making this test system suitable for the detection of possible consequences of RF-EMF exposure. Important biological endpoints such as fertility, reproduction, behavior and development are rather easy to implement, especially in a laboratory setting.

Besides the fruit fly as model organism, special ecological relevance is outlined by pollinators, in particular by the honey bee *Apis mellifera*. They provide critical pollination services valued at over $200 billion worldwide (Lautenbach et al. 2012), representing 9.5 % of the total human food production (Gallai et al. 2009). However, bees have suffered periodic losses within the last century, and in the US a phenomenon called colony collapse disorder (CCD) made headlines in the first decade of the new millennium (vanEngelsdorp et al. 2009). Several causative factors have been outlined in the past, among others, pathogens, malnutrition, management, and pesticides have been narrowly focused as main culprits (Steinhauer et al. 2018).

Many other factors were also considered to have an impact on honey bee health, however with a rather insignificant regard. A few to name are air pollution (Girling et al., 2013; McFrederick et al., 2008), nanomaterials (Milivojevic et al., 2015), solar radiation (Ferrari, 2014), robbing insects (Core et al., 2012) and global warming (Le Conte & Navajas, 2008). Worthy of mention, in 2007 a story in an UK newspaper brought to the fore that CCD can be linked to RF-EMF with drastic consequences for bee behavior and homing success (Kimmel et al. 2007; Carreck, 2014). Subsequent studies seem to provide supporting evidence of impaired behavior (Favre, 2011) and affected homing ability (Ferrari, 2014), bearing a potential risk to other bee species such as bumblebees (*Bombus terrestris*), when interacting with floral electric fields and electric field sensing as important sensory modality (Clarke et al., 2013).

However, there are far too few scientific publications to draw a clear conclusion in regard if and to which extent mobile phone radiation represents a real threat to honey bees. A current review actually goes as far as stating that all examined studies were characterized by substantial shortcomings which were sometimes even admitted by their authors upfront (Verschaeve, 2014).

For a honey bee colony, health and productivity is directly linked to its queen. She represents the growth potential expressed as productivity, being the only egg layer in the collective and therefore responsible for a positive turnover of workers to increase in size at the beginning of each bee season (Moore et al. 2015). In an US survey of winter colony losses, the fourth most important factor identified was due to queen failure (vanEngelsdorp et al., 2008). Given the importance of this individual, our experiments therefore strictly focused on ontogenetic development and further mating success of young queens. We have created a worst case scenario, where mobile phone radiation was adopted by natural means of human exposure. To our knowledge this is the first study that analyzes the effect of a chronic application of mobile phone radiation on honey bee queens. We wanted to prove (i) if under field conditions and good apicultural practice the radiation has any effect at all and to what extent, in addition (ii) we wanted to follow queens which developed under chronic RF-EMF exposure to assess potential risks for the bee colony.

## 2 MATERIALS & METHODS

### 2.1 Field sites and weather conditions

The field sites were located near the Apicultural State Institute in Stuttgart-Hohenheim, Southern Germany (48°42’31.8”N 9°12’38.2”E). At the time present, natural food sources consisted mainly of nectar from diverse local flora such as *Taraxacum officinale*, *Rubus section*, *Tilia spp*. and others. The average temperature during the experiment ranged from 15.2 to 20.1 °C with a precipitation of 90 to 45 L/m². Overall, good weather conditions prevailed for both, mating and foraging (DWD 2018).

### 2.2 Experimental setup

This study was performed from May until August in 2018 with healthy queenright colonies from the stock of our apiary. Two replications were employed simultaneously, consisting of two collector colonies: Rep1 (Control1 + EMF1) and Rep2= (Control2 + EMF2). For both approaches, one brood frame with almost fully covered areas of sealed brood and attached bees from eight random colonies were taken out on D-9 and placed in a new ten-frame box, respectively. This box was supplied with two frames of food, as well as a second box on top with ten food frames to ensure sustenance and sufficient room for the hatching bees. Nine days after this procedure (D0), the hive was inspected, and where appropriate, supersedure cells were removed to prevent the introduction of a young queen. Further, 18 frames then were split homogeneously but random into two boxes with nine frames each, complemented with a grafting frame in the center. L1 larvae from a selected colony were grafted and introduced, respectively. Again, grafting of the larvae was randomized by using both sides of the brood comb (A and B). Per replication, 26 larvae (13 A, 13 B) were assigned to each treatment, i.e. control and EMF.

The two boxes then were placed at a different location in approximately 3 km distance to prevent worker bees to return to their original position. Subsequently, at different intervals, assessments were performed to check the no. of accepted larvae after grafting (D1), to protect the capped cells before hatching (D10), to check the hatching rate (D13) and the mating success (D24). After the young queens have hatched, they were transferred to mating units consisting of one of the former brood frames with approximately 1,000 bees attached and one food comb.

Successful mating was confirmed on D24 by the presence of eggs, young larvae and capped brood and queens from each treatment (five from the control, four from the treatment) were re-accommodated in new 10-frame boxes to develop into full colonies. After approximately twelve weeks (D88), a colony assessment was performed to record the number of bees and brood. See Fig. 1 for a detailed timeline.

**Fig. 1.**
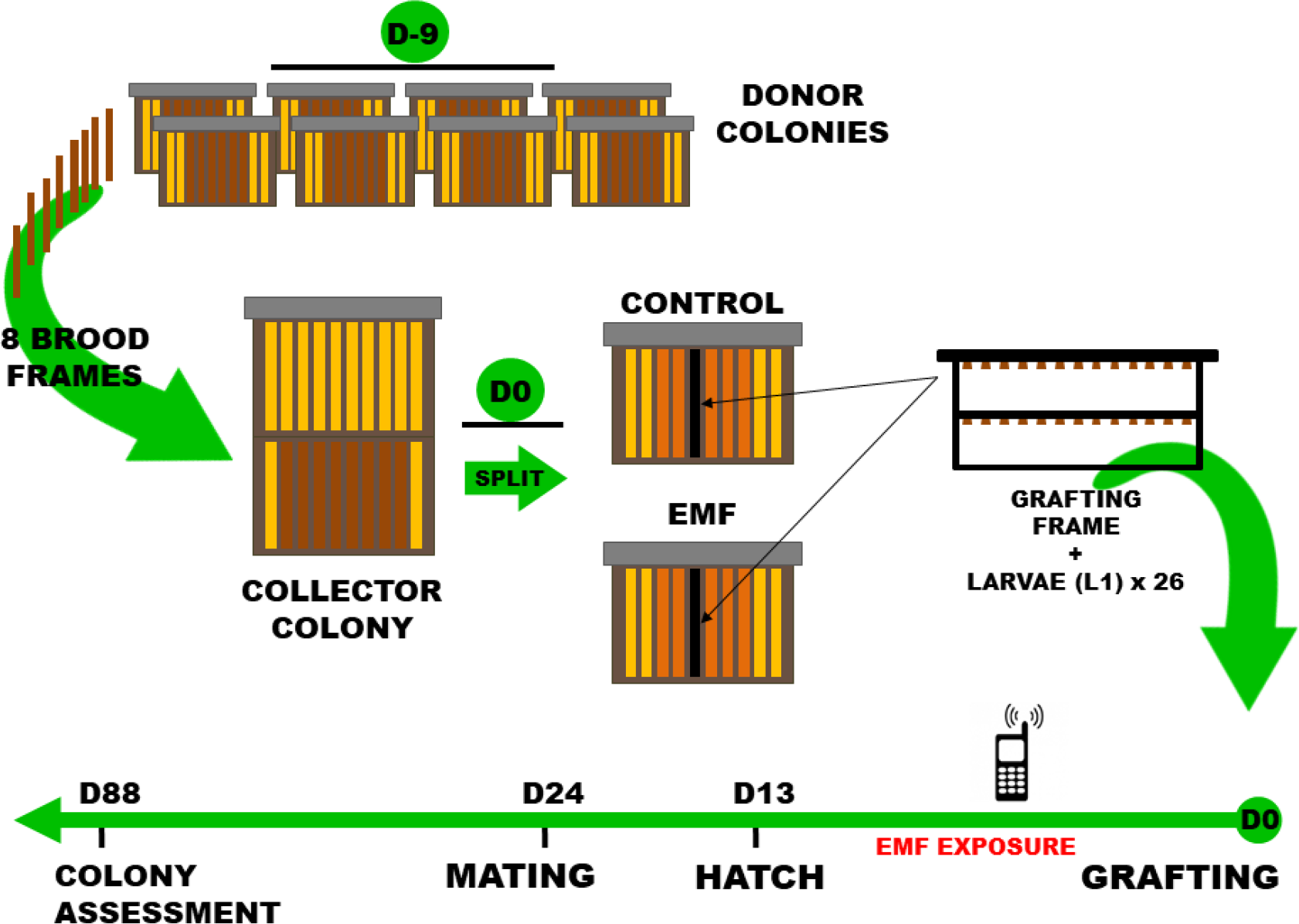
Timeline of the experiment. At D-9, eight brood frames with attached bees were taken out from respective donor colonies and placed in one collector colony. At D0, supersedure cells were removed and the collector colony was split in two sub-colonies. In addition, a grafting frame with L1 larvae was inserted. RF-EMF exposure lasted until D13, when queens were about to hatch. Young queens were subsequently inserted into mating units where mating success was checked at D24. Successfully mated queens with one frame of approximately 1,000 bees were relocated into new boxes where they were able to establish a new colony. Finally, at D88 the condition of these colonies was assessed.

### 2.3 EMF treatment

Queen larvae/pupae were treated with a mobile phone (AEG M1220, GSM quad band: 800/900/1800/1900 MHz, China) attached to the grafting frame holding 26 queen cups (Nicot, NICOTPLAST SAS, Maisod, France), this device was turned off in the control group for sham exposure. To ensure power supply, the phone was equipped with a power bank (PLOCHY 24,000 mAh Solar, China), the battery status was frequently checked. After the larvae were grafted into the cups by using an appropriate tool, 15 telephone calls with a two minute duration were applied daily for a total of two weeks (non-modulated emission) at random. The radiation was measured three times in three different distances to the mobile phone with a fixed instrument illustrated in Fig. 2 (Pyle PMD74, Calibration: 2450 MHz, measurement range: 0-15 mW/cm^2^, China) to verify an adequate EMF output.

**Fig. 2.**
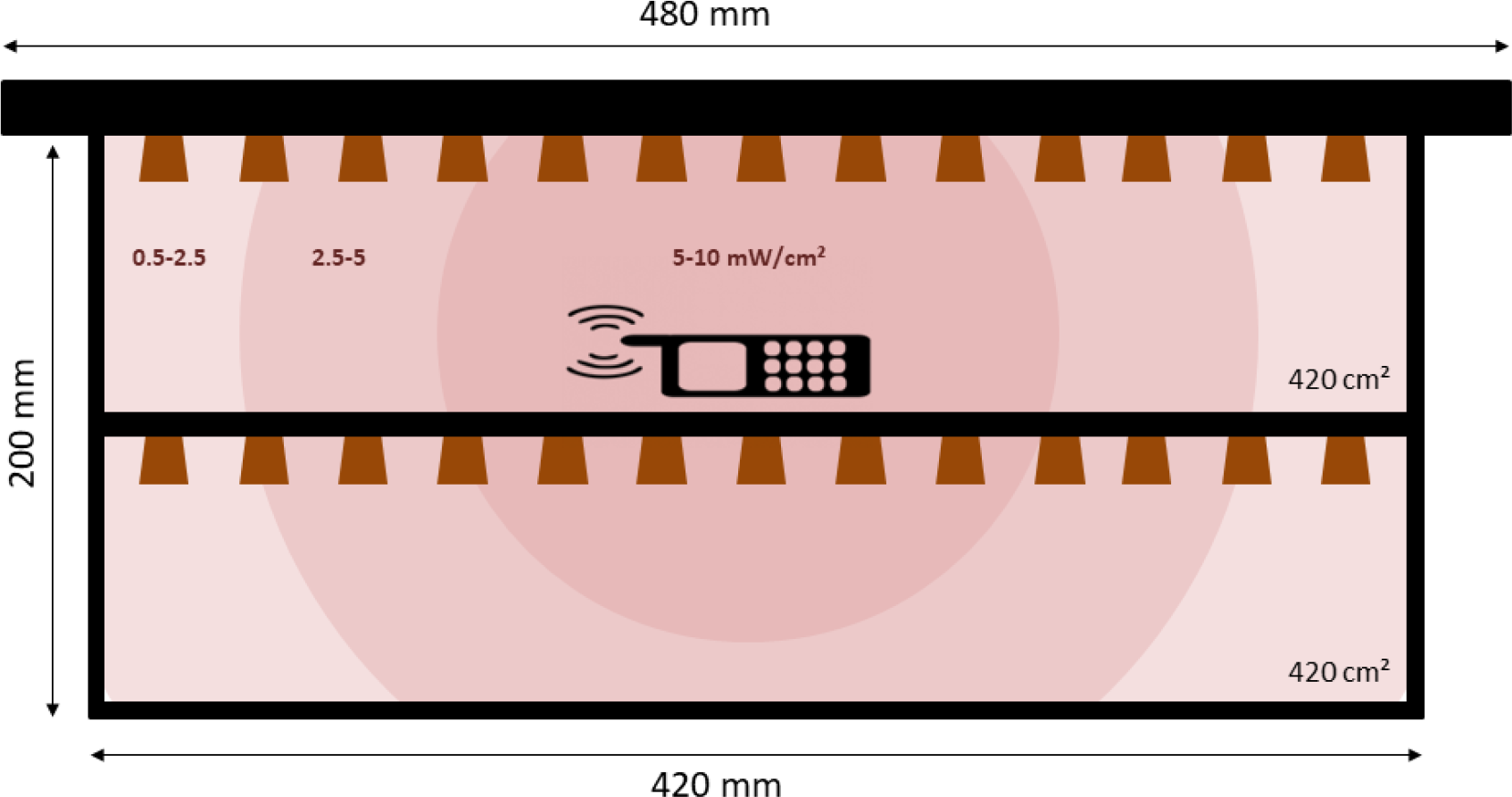
Grafting frame placed in the EMF treatment colony containing 26 queen cups. The mobile phone device was attached in the center of the frame, its radiation intensity is indicated with the differently colored sections in the illustration (dark > light)

### 2.4 Colony assessment

The amount of bees and brood cells (open and sealed) were estimated with the Liebefeld Method (Imdorf et al., 1987), which is a feasible tool to provide accurate and reliable evaluation of colony strength (measuring error +/− 10 %). Care was taken that all colonies were evaluated by the same person to minimize variation and colony assessment was conducted in the morning before bee flight.

### 2.5 Statistical analysis

We evaluated the mortality data with a Kaplan-Meier-Survival analysis. Survivorship between control and treatment was compared pairwise and tested for significance with a Log-Rank Tests (Cox-Mantel). Individuals collected at the end of the experiment were considered censored, as were those observed but not collected on the final day. Furthermore, larvae that disappeared during the experiment were considered dead on the last day they were seen. Both treatment groups and the two replicates (Rep1= Control1 + EMF1; Rep2= Control2 + EMF2) were additionally compared with a Cox proportional hazards model to determine the hazard ratio (HR). Possible inter-colony effects were evaluated as covariate to justify pooling data of the same treatments. The estimated number of bees and brood cells were checked with a Shapiro-Wilk test for normal distribution. If data was normal, a one-way ANOVA was performed on the two experimental groups, respectively. For all tests RStudio (R Core Team, 2018) and significance level of α=0.05 was used.

## 3 RESULTS

### 3.1 Honey bee queen survival

The Kaplan-Meier-Survival analysis of both groups showed a significant difference indicating a higher mortality of the EMF treated bees when compared to the control group (p=0.0054) (Fig. 3). In addition, a Cox proportional hazards model was applied to determine the hazard ratio (HR) displayed as forest plot (Fig. 4). With a HR of 2.3 the EMF treated queens had a significantly increased risk of dying when compared to the control (p=0.003). Moreover, the two replicates (Rep1 and Rep2) were compared as covariate to display possible inter-colony effects. However, with a HR of 1.7 queens in Rep2 did not have a higher risk of dying when compared to Rep1 (p=0.062), therefore data of both replicates were pooled.

**Fig. 3.**
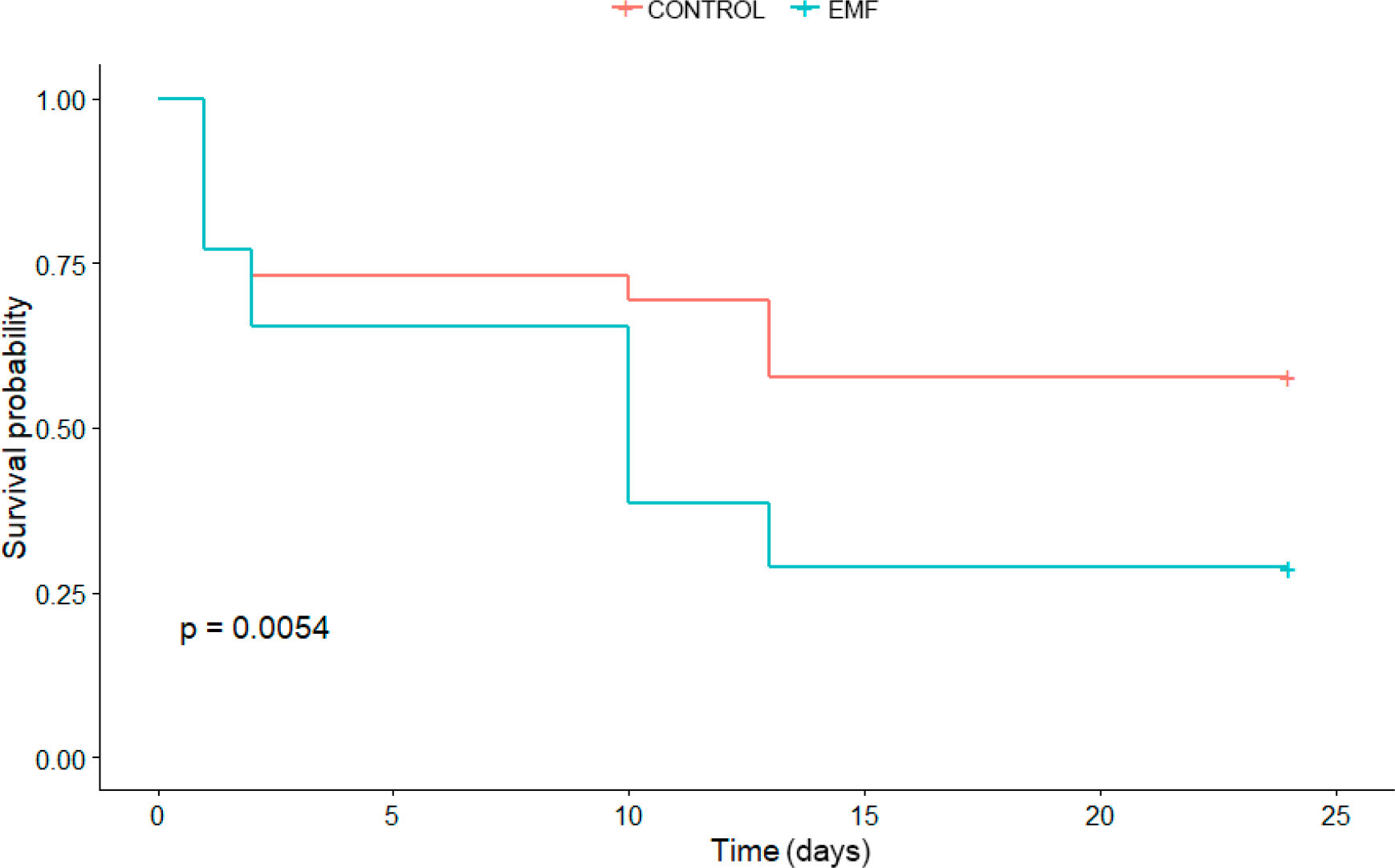
Both groups were compared with a Kaplan-Meier-Survival analysis. A post-hoc Log-Rank test (Cox-Mantel) revealed a significant higher mortality in the EMF treatment when compared to the control (Log-Rank p=0.0054), where a significant decrease of individuals occurred during the pupation phase of the experiment (see also Fig. 5)

**Fig. 4.**
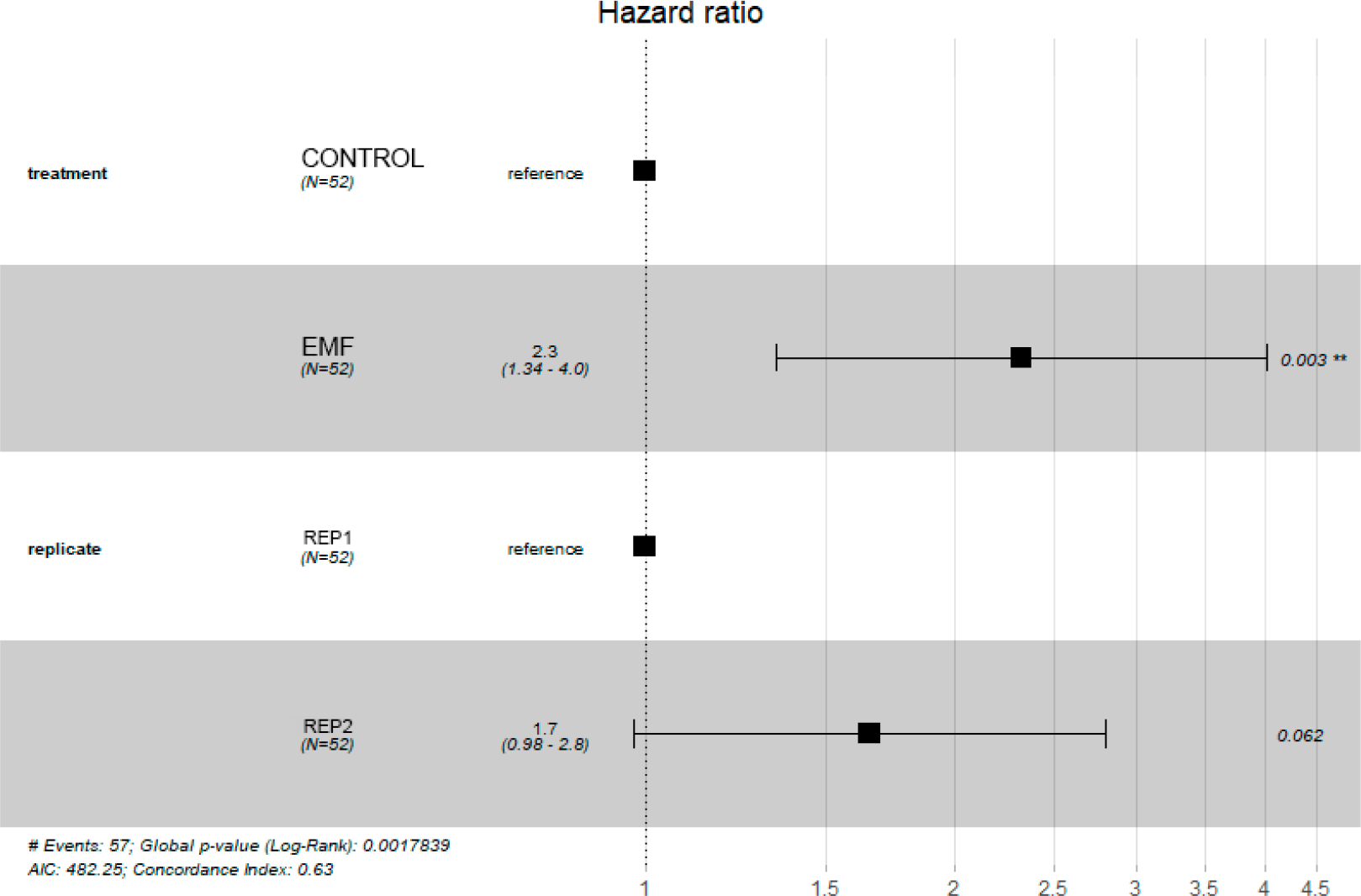
Both treatment groups and the two replicates (Rep1 and Rep2) were additionally compared with a Cox proportional hazards model to determine the hazard ratio (HR) displayed as forest plot. With a HR of 2.3 the EMF treated queens had a significantly increased risk of dying when compared to the control (p=0.003). And with a HR of 1.7 queens in Rep2 did not have a higher risk of dying when compared to Rep1 (p=0.062)

### 3.2 Hatching and mating success

The acceptance rate of grafted larvae on D1 was 76.9 % and identical in both treatments. As shown in Fig. 3, a significant decrease of individuals in the EMF treatment occurred during the pupation phase of the experiment. At D10, queen cells were protected with a cage to prevent hatching queens from killing each other. We observed a ratio of 73.1 (control) to 65.4 % (treatment) at this stage compared to the initially grafted cells. The hatch of adult queens at D13 revealed a significant decrease of formerly treated queens during pupation with a proportion of 69.2 % in the control to 38.5 % in the treatment, levelling out with a similar decrease of both groups at the assessment of mating success at D24 (control 57.7 %, treatment 28.8 %) (for statistical evaluations see also Fig. 3).

### 3.3 Colony assessment

The population of bees and brood cells was estimated at D88. The results are shown in Fig. 6A for the number of bees and in Fig. 6B for the number of brood cells. We compared the two treatment groups with a one-way ANOVA but could not see significant differences for the number of bees (p=0.688) or the amount of brood cells (p=0.768).

**Fig. 5.**
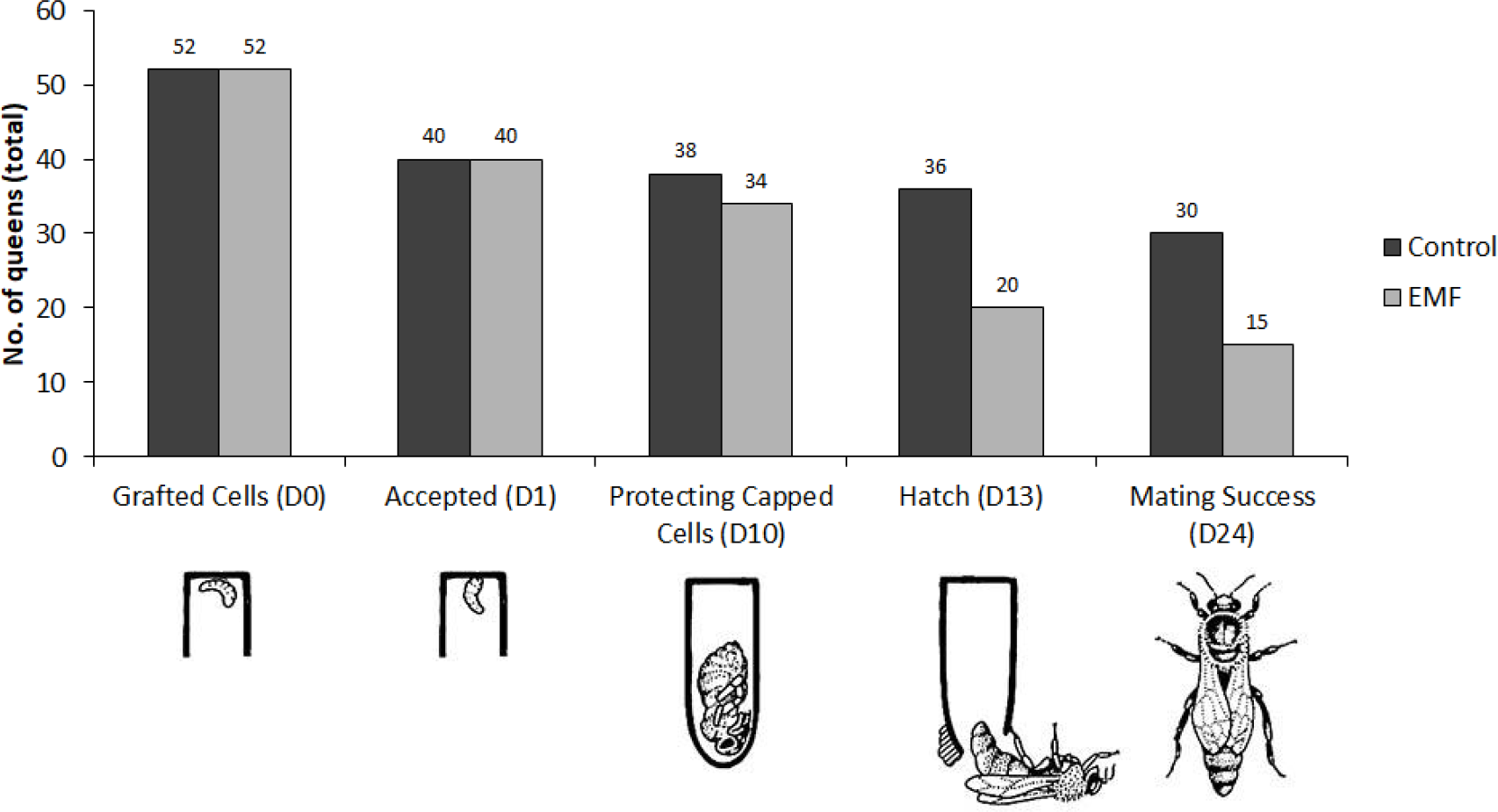
Total number of L1 larvae grafted and followed through their ontogenetic development from pupa to adult. In the EMF treatment a significant decrease of individuals came into effect within the pupation phase of the experiment (see also Fig. 3). Illustrations after Gullan & Cranston 2014

**Fig. 6A.**
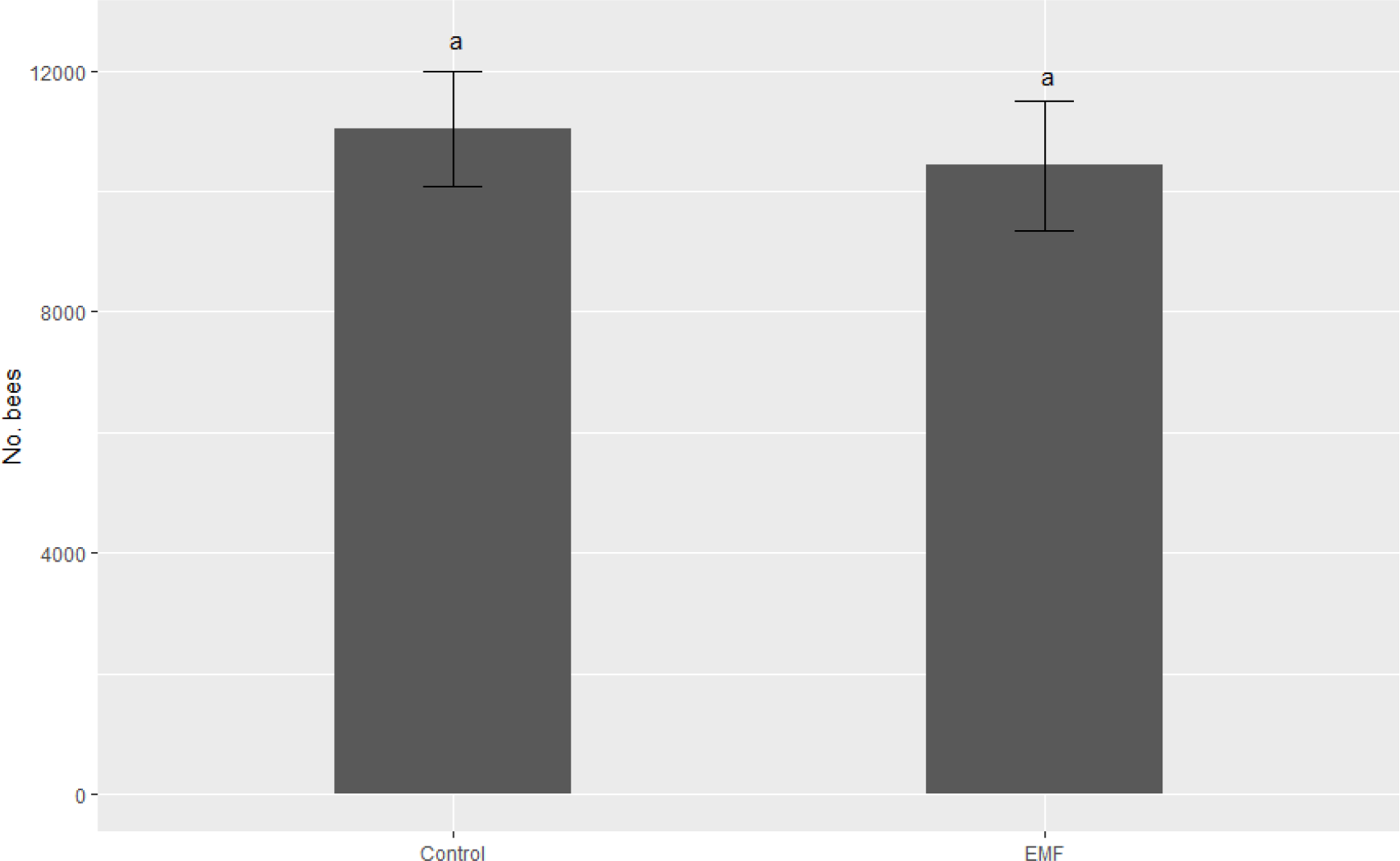
Number of bees estimated at D88 in the colonies of the control (n=5) and of the EMF treatment (n=4). Same letters indicate no statistically significantly differences (p=0.688, ANOVA).

**Fig. 6B.**
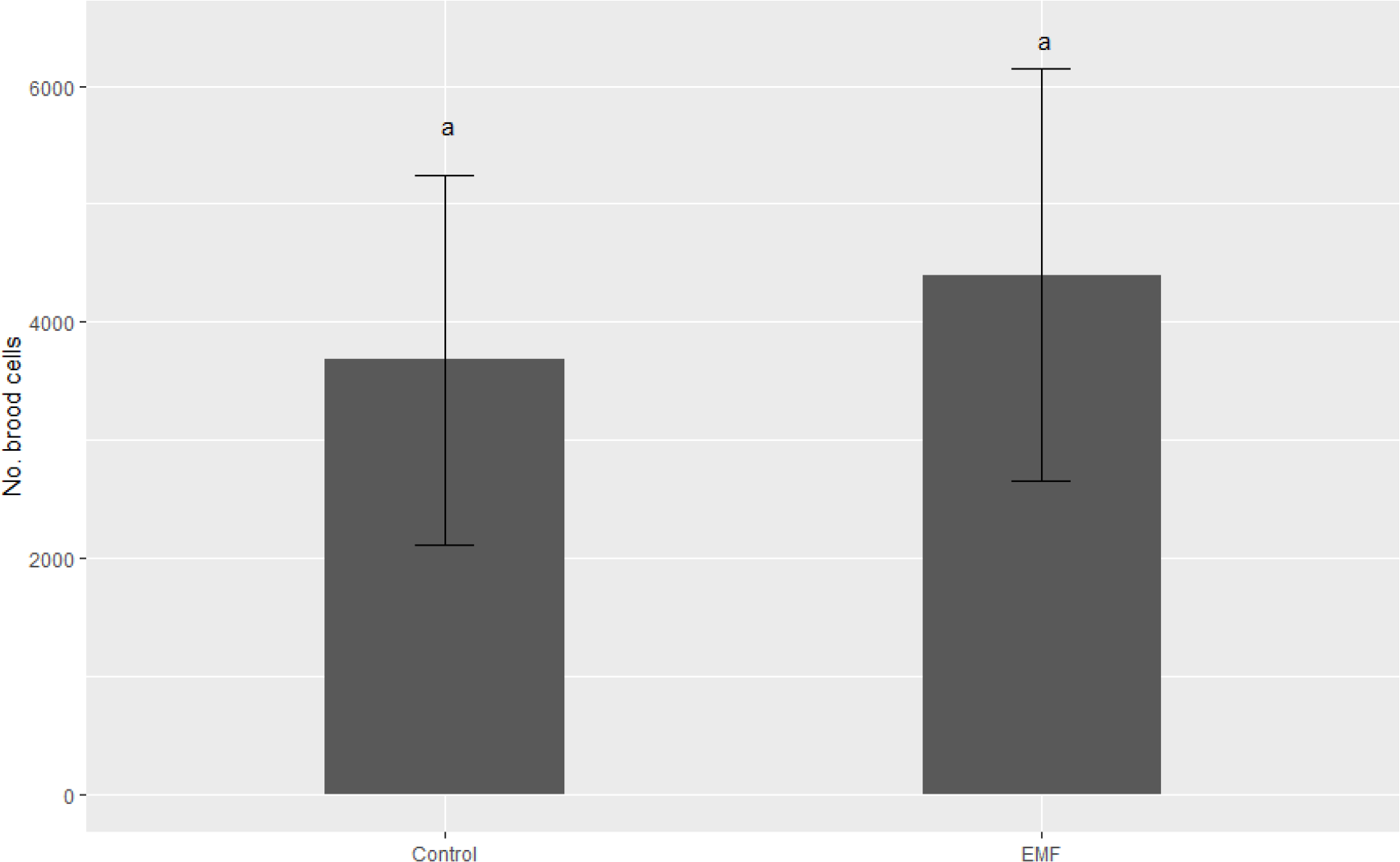
Number of brood cells estimated at D88 in the colonies of the control (n=5) and of the EMF treatment (n=4). Same letters indicate no statistically significantly differences (p=0.768, ANOVA).

## 4 DISCUSSION

The emission of radiofrequency electromagnetic fields (RF-EMF) and their negative effects towards honey bee health has been controversially discussed in the past (Carreck 2014; Verschaeve 2014; Panagopoulos et al. 2016). Here, we could demonstrate for the first time that RF-EMF exposure has significantly affected ontogenetic queen development under field conditions. We observed an increased mortality during pupation resulting in a reduced hatching rate of the later queens. This is in line with a reduced reproductive capacity found in fruit flies (*Drosophila melanogaster*) (Panagopoulos et al. 2004, Margaritis et al. 2014), where a linear decrease of fecundity was reported with the frequency of exposure (Panagopoulos and Margaritis 2010). This decrease was further associated with the distance to the mobile phone device showing the strongest effects at < 10 cm (Panagopoulos et al. 2010). In our setup, the most distant queen cups were approximately 21 cm away from the radiation source and we therefore assume that for all larvae a worst case scenario came into effect. In addition, the impairment of fruit flies seemingly depended on field intensity (Panagopoulos et al. 2007) not only reducing the offspring but also the ovarian size of the exposed subjects (Panagopoulos 2012).

At present, only a few studies have investigated the influence of irradiation on insect development. As an example, larvae and pupae of the dried fruit beetle (*Carpophilus hemipterus*) and the yellow fever mosquito (*Aedes aegypti*) were exposed to Gamma radiation (ionizing radiation). The radiotherapy did not cause acute death in larvae but decreased pupation significantly, no effects however could be observed when either young or old pupae were exposed (Johnson 1987, Akter & Khan 2014). It seems likely that RF-EMF had a similar effect in our study, as larval mortality was not elevated. However, this should be further underpinned by exposing larvae and pupae separately. Moreover, Vilić et al. (2017) found honey bee worker larvae significantly affected when exposed to modulated but not to non-modulated RF-EMF radiation, resulting in DNA damage and further corroborating our hypothesis as we only have used non-modulated fields.

In addition, we could show that mating success remained unaffected suggesting that navigation and the possible disruption of magnetoreception came not into effect or was at least not long-lasting (Vácha et al. 2009). Interestingly, we provide evidence that developing honey bee queens once they have survived RF-EMF exposure seem to retain the ability to establish an intact colony. This is indicated by similarly strong numbers of bees and the amount of brood in both our treatment groups with the absence of any signs of impairment (e.g. patchy brood pattern). As a further critical step of colony survival however, overwintering should also be assessed to elucidate possible long term effects from the irradiation (Smart et al. 2016).

The social entity as a whole is able to buffer environmental stressor of various kinds as an expression of social resilience (Straub et al. 2015). Worker bees are nursing eggs and feeding larvae of different casts in their social state, potentially contributing to this mechanism. Here we focused on the development of individual queens from larvae to adult, however, the outcome of our study could also be influenced by the condition of the collector colonies that we have created but not further assessed. Eggs, larvae and pupae are very sensitive stages of development and intensive care is taken to supply their substantial needs in terms of nutrition and environmental conditions, i.e. maintaining a constant temperature and humidity (Wang et al. 2015, Eouzan et al. 2018). RF-EMF radiation is known to affect bees behavior in different ways (Favre 2011, Ferrari 2014), which makes it plausible that brood care could also be adversely affected. This important factor should be further investigated and included in future experiments.

With an increasing number of mobile phone devices and as a consequence of good accessibility a higher density of phone masts, not only urban but also rural areas in particular are more and more exposed to irradiation (Balmori 2009). A measurement of RF-EMF intensities across different European cities revealed maximum radiation values ranging from 0.84 to 0.59 V/m corresponding to 92.33 and 187.16 nW/cm² (Urbinello et al. 2014a), respectively, with a maximum value of 127 nW/cm² in public transport (Sagar et al. 2016). In contrast, the power flux density measured in our study seemed to be way beyond these values, demonstrating that the intermittent stress on the test subject(s) can be many fold higher than average levels measured in the surroundings, emitted from generators or found in agglomerations. Our findings confirm that there is a high variability in mobile phone emission (Frei et al. 2009), representing an important feature in terms of bioactivity towards living organism’s defense against environmental stressors (Panagopoulos et al. 2015). The authors therefore suggest not using simulated but real mobile phone emissions in an experimental setup, which we have considered. In addition, we have tried to apply a human exposure scenario in terms of average number of calls and average call duration performed with mobile phone devices. The mobile call duration reported by the German Federal Network Agency (2011) was 2.5 min per day, in Shum et al. (2011) ranging from 2.1 min (self-reports) to 2.8 min (billing records) and < 2 min in Friebel & Seabright (2011). Further, the average number of calls per day ranging from 4.1 (Shum et al. 2011) to 5 per day in adults (Lenhart 2010). In contrast, an average of 33.1 min was reported for total mobile phone call duration from undergraduate college students per day in the US (Roberts et al. 2014). We therefore decided to employ 2 min per call and 15 calls per day resulting in 30 min exposure per day in our experiment, representing a realistic human exposure.

Different exposure scenarios were applied in honey bees and a broad range of effects are reported (Cucurachi et al. 2013). Some studies even claimed with RF-EMF to have found the major cause for CCD (Carreck 2014). However, many of these studies had substantial deficits such as a very low sample size (Sharma & Kumar 2010), intransparent methods (Sahib 2011, Kumar et al. 2011, Dalio 2015) or were even preliminary and did not undergo peer-review (Kimmel et al. 2007). Therefore, findings of this quality were generally not considered reliable in their contribution to colony losses and are far from conclusive (Carreck 2014). To achieve a broader understanding how RF-EMF potentially influences the honey bee superorganism, it is mandatory to emphasize the conditions under which the study was conducted, particularly the level and duration of exposure, in the presence of the relevant environmental situation (Verschaeve 2014).

As a trend of the last decades, beekeeping became famous with the life style of townsmen all across the globe (Lorenz & Stark 2015, Kohsaka et al. 2017, Stange et al. 2017). Therefore, density of bee colonies held in urban areas has dramatically increased and may favor the spread of diseases or pathogens (Youngsteadt et al. 2015). However, following this trend also bears the risk of a higher exposure to RF-EMF emission, which seems to be continuously increasing in major cities (Urbinello et al. 2014b), potentially affecting bee health in a future scenario. It might also be worthy to look into parasite-host-interactions of the honey bee, *Varroa destructor* in particular, where a disturbance through RF-EMF in host-finding could actually be a benefit (Frey et al. 2013). Surprisingly, not many studies are available that are investigating the influence of such irradiation on bees and other important pollinators. It has even been suggested to create pollinator reservoirs beneath power corridors for an optimal land use and as a benefit for many insects (Russel et al. 2018). Yet, it remains unclear to what extend electromagnetic fields can possibly influence these microenvironments.

## Conclusion

Even though detrimental effects on ontogenetic queen development were revealed by the outcome of our study, caution is needed in interpreting these results. We have created by far a worst case scenario to which honey bee colonies would not be exposed under realistic conditions. Duration and level were similar to average human exposure by the use of a mobile phone, but not to those present at an apiary, neither in rural nor in urban areas. And yet, queens that survived the treatment were able to establish full functional colonies, demonstrating an immense recovering potential. Therefore we do not assume any acute negative effects on bee health in the mid-term. However, we do not rule out an influence through lower doses of permanent irradiation, in particular on a chronic sublethal level. Hence, we urgently suggest further research should be carried out in the long-term to ascertain what impacts are to be expected.

## ACKNOWLEDGEMENTS

Many thanks to Simon Hummel for providing adequate breeding lines to conduct research with extremely calm honey bees.

This research did not receive any specific grant from funding agencies in the public, commercial, or not-for-profit sectors.

Declarations of interest: none.

Ethical approval: This article does not contain any studies with human participants or animals performed by any of the authors.

